# The role of chloroplast SRP54 domains and its C-terminal tail region in post- and cotranslational protein transport *in vivo*

**DOI:** 10.1101/2024.04.21.590438

**Authors:** Annika Bischoff, Jennifer Ortelt, Beatrix Dünschede, Victor Zegarra, Patricia Bedrunka, Gert Bange, Danja Schünemann

**Author notes:** Correspondence: Prof. Dr. Danja Schünemann.

## Abstract

In the chloroplast, the 54 kDa subunit of the signal recognition particle (cpSRP54) is involved in the posttranslational transport of the light-harvesting chlorophyll a/b-binding proteins (LHCPs) and the cotranslational transport of plastid-encoded subunits of the photosynthetic complexes to the thylakoid membrane. It forms a high-affinity complex with plastid-specific cpSRP43 for posttranslational transport, while a ribosome-associated pool coordinates its cotranslational function. CpSRP54 constitutes a conserved multidomain protein, comprising a GTPase (NG) and a methionine-rich (M) domain linked by a flexible region. It is further characterized by a plastid-specific C-terminal tail region containing the cpSRP43-binding motif. To characterize the physiological role of the various regions of cpSRP54 in thylakoid membrane protein transport, we generated *Arabidopsis thaliana* cpSRP54 knockout (*ffc1-2*) lines producing truncated cpSRP54 variants or a GTPase point mutation variant. Phenotypic characterization of the complementation lines demonstrated that the C-terminal tail region of cpSRP54 plays an important role specifically in posttranslational LHCP transport. Furthermore, we show that the GTPase activity of cpSRP54 plays an essential role in the transport pathways for both nuclear-as well as plastid-encoded proteins. In addition, our data revealed that plants expressing cpSRP54 without the C-terminal region exhibit a strongly increased accumulation of a photosystem I assembly intermediate.

**Highlight:** This study elucidates the important role of the chloroplast SRP subunit, cpSRP54, in the biogenesis of both the LHC antenna and the cores of photosystem I and II *in vivo*.

## Introduction

The light-harvesting chlorophyll a/b-binding proteins (LHCPs) are the most abundant integral proteins of the thylakoid membrane and play a pivotal role in capturing and channeling solar energy to the photosystem (PS) I and II reaction centers. In *Arabidopsis thaliana*, the PSII antenna (LHCII) is mainly formed by the LHCP family members Lhcb1-6, while the proteins Lhca1-4 are the major constituents of the PSI antenna (LHCI) (Ben-Shem et al., 2003; Nelson and Ben-Shem, 2004; Mazor et al., 2017; van Bezouwen et al., 2017). The biogenesis of the nuclear-encoded LHCPs requires a series of orchestrated steps to ensure the proper targeting and integration into the thylakoid membrane.

After import of the LHCP through the TOC/TIC translocation machinery of the chloroplast envelope it is bound by the posttranslationally acting stromal chloroplast signal recognition particle (cpSRP) (Akopian et al., 2013; Ziehe et al., 2018). CpSRP is a high affinity heterodimer formed by the 43 kDa and 54 kDa subunits cpSRP43 and cpSRP54, respectively (Franklin and Hoffman, 1993; Schünemann et al., 1998; Klimyuk et al., 1999; Gao et al., 2015; Ziehe et al., 2018). The LHCP/cpSRP complex (transit complex) docks to the thylakoid membrane through an interaction with the cpSRP54 receptor, cpFtsY, and the Alb3 insertase (Kogata et al., 1999; Tu et al., 1999; Moore et al., 2000; Asakura et al., 2008). *In vitro* experiments demonstrated that the insertion of LHCP into the membrane requires GTP (Hoffman and Franklin, 1994; Yuan et al., 2002). GTP triggers the GTPase cycle of the cpSRP54/cpFtsY complex as both proteins interact directly by their GTPase coding NG-domains, which stimulates each other’s hydrolysis activity and dissociation of the complex (Jaru-Ampornpan et al., 2007; Jaru-Ampornpan et al., 2009). A central step in LHCP transport is the accommodation of its hydrophobic domains by cpSRP to maintain its solubility and insertion competence. Here, cpSRP43 plays a key role, as it interacts directly with a conserved DPLG motif between the second and third transmembrane helix of LHCP and is sufficient to prevent LHCP from aggregation (DeLille et al., 2000; Tu et al., 2000; Jaru-Ampornpan et al., 2010; Falk and Sinning, 2010). Furthermore, it has been suggested that LHCPs can be sorted to the thylakoid membrane by cpSRP43 alone at least in a *A. thaliana* double mutant lacking cpSRP54 and cpFtsY (Tzvetkova-Chevolleau et al., 2007). The role of cpSRP54 in transit complex formation is less defined. While some data indicate that cpSRP54 plays an indirect role in transit complex formation by driving cpSRP43 into an active state that allows tight binding to LHCPs, other data point to a direct binding of cpSRP54 to the third transmembrane helix of LHCP (High et al., 1997; Groves et al., 2001; Henderson et al., 2016; Liang et al., 2016). The interaction between cpSRP54 and cpSRP43 has been extensively investigated, highlighting its dependence on the positively charged “ARRKR” motif situated in the C-terminal tail region of cpSRP54. This motif facilitates binding to the second chromodomain of cpSRP43 (Funke et al., 2005; Holdermann et al., 2012; Dünschede et al., 2015).

Remarkably, a secondary pool of stromal cpSRP54 is linked with chloroplast ribosomes, playing a crucial role in facilitating the efficient cotranslational sorting of plastid-encoded multi-span thylakoid membrane proteins. These include components like the reaction center subunits of Photosystem I (PsaA, PsaB) and Photosystem II (PsbA, PsbD) (Franklin and Hoffman, 1993; Schünemann et al., 1998; Hristou et al., 2019). The ribosomal binding interface on cpSRP54 has been assigned to two binding motifs, one corresponding to the ARRKR motif in the C-terminal tail and the second formed by a short motif within the M-domain of cpSRP54 (Hristou et al., 2019) (Figure 1A). Furthermore, a direct contact of cpSRP54 with a hydrophobic region of the nascent PsbA chain was demonstrated (Nilsson et al., 1999; Nilsson and van Wijk, 2002). While the precise molecular mechanism governing the targeting and insertion of nascent chains into the thylakoid membrane remains largely unresolved, current data indicate that a contact between cpSRP54 and cpFtsY facilitates the docking of the translating ribosome to the thylakoid membrane. Subsequently, membrane insertion is thought to be mediated by the cpSec1/Alb3 insertion machinery (Zhang et al., 2001; Göhre et al., 2006; Tzvetkova-Chevolleau et al., 2007; Walter et al., 2015a; Walter et al., 2015b).

**Figure 1:**
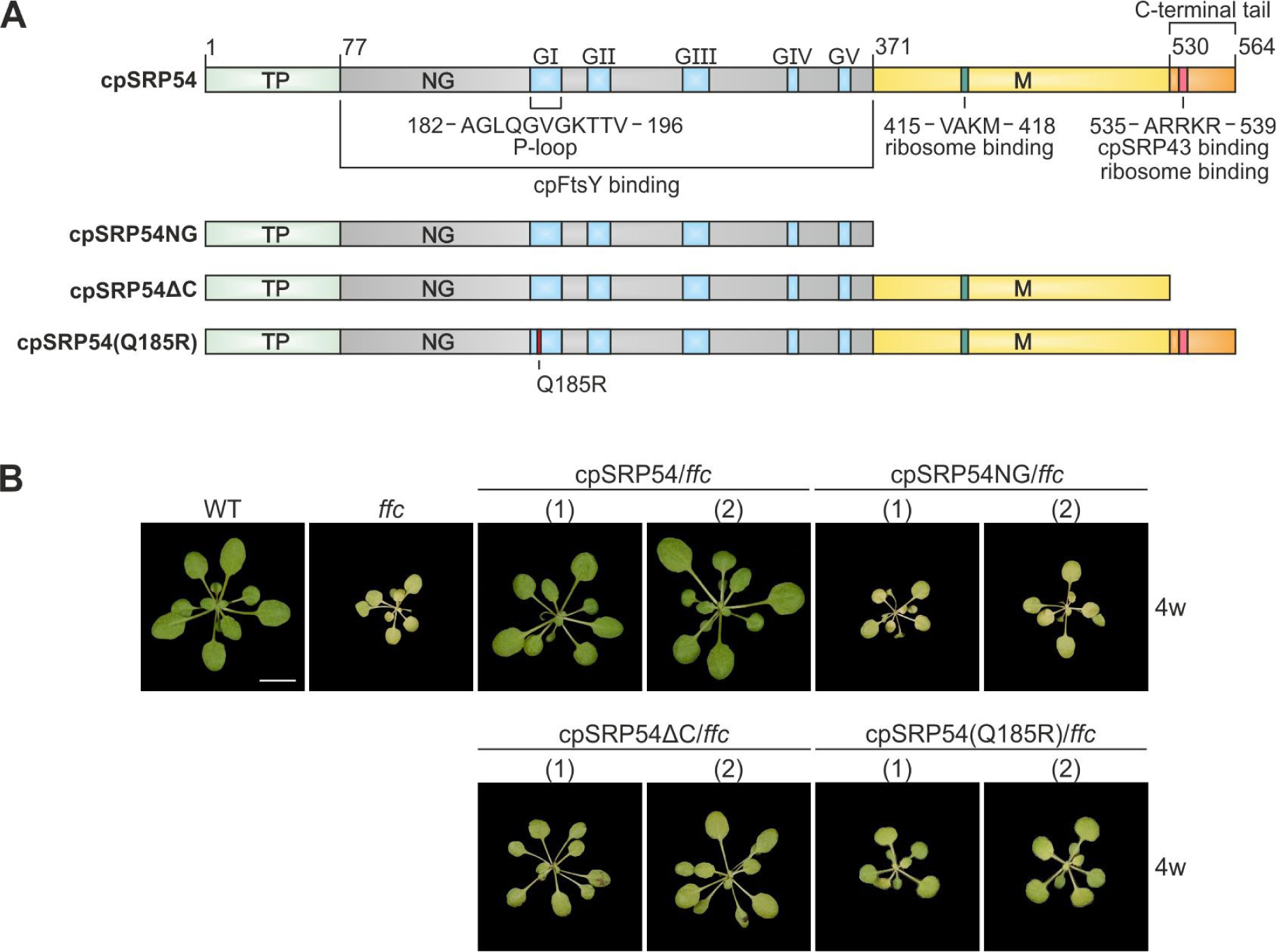
*Arabidopsis thaliana* cpSRP54 variants and complementation lines of *ffc*. (**A**) Scheme of the *A. thaliana* cpSRP54 protein and variants used to generate *ffc*-complementation lines. The multidomain protein cpSRP54 is composed of an amino-terminal N-domain in close contact to a G-domain (NG), a methionine-rich M-domain and a C-terminal tail region. The G-domain provides the interaction region of cpSRP54 and its receptor cpFtsY including the conserved GTPase motifs GI (phosphate (P)-binding loop) – GV. The cpSRP43 binding motif, which serves also as one of the two ribosomal binding sites is located within the C-terminal tail region. The second ribosomal binding motif is located within the M-domain. Scheme according to Ziehe et al., 2018. (**B**) Phenotypes of 4-week-old *A. thaliana ffc*-complementation lines compared with wild type (WT) and the cpSRP54-lacking mutant *ffc*. The scale (white line) corresponds to 1 cm.

This study was performed to analyze the physiological role of the individual domains of cpSRP54 in post– and cotranslational protein sorting *in vivo*. We transformed *ffc1-2 A. thaliana* plants (lacking cpSRP54) with cpSRP54 constructs encoding C-terminal truncations or the point mutation variant cpSRP54(Q185R), which is impaired in GTP hydrolysis activity. Our data show that the plant-specific C-terminal tail region of cpSRP54 is required for posttranslational transport of the LHCPs *in vivo*. In addition, we provide *in vivo* evidence that the GTPase activity of cpSRP54 is essential for posttranslational LHCP sorting as well as for cotranslational insertion. Our study also shows the accumulation of a PSI assembly intermediate in the *ffc1-2* line expressing a cpSRP54 variant, which lacks its C-terminal extension. Furthermore, our data indicate that the assembly of LHCI to PSI is required for the incorporation of PsaK into PSI.

## Materials and methods

### Generation of Arabidopsis thaliana complementation lines

To generate complementation lines, *A. thaliana ffc* mutant plants (*ffc* 1-2, previously described in (Amin et al., 1999)) were transformed with the plant transformation vector pCBi18 (Biesgen and Weiler, 1999; Lehmann et al., 2017) containing cDNA fragments of *A. thaliana* cpSRP54 (Accession number: At5g03940) variants. The full-length coding sequence of *A. thaliana* cpSRP54 was obtained from the Arabidopsis biological resource center (ABRC Stock Number: U10271). The residues 1-564 (At-cpSRP54), 1-370 (At-cpSRP54NG) and 530-564 (At-cpSRP54ΔC) were introduced downstream the CaMV 35S promoter into pCBi18, replacing the originally included uidA gene via XbaI/SacI restriction sites. For plasmid construction, gene-specific primers adding a His-tag sequence were used with the In-Fusion® HD EcoDry™ kit (Takara). The complementation construct containing *A. thaliana* cpSRP54 residues 1-564 including a point mutation at position 185 (At-cpSRP54(Q185R)) was generated using the QuikChange Lightning site-directed mutagenesis kit (Agilent Technologies) using pCBi18-At-cpSRP54 as template. Used primers are listed in Supplementary Table 1. All plasmids were checked by Sanger sequencing and used for *Agrobacterium tumefaciens*-mediated plant transformation (Block, 1993; Chateau et al., 2000). The plasmids were introduced into *Agrobacterium tumefaciens* GV3101 by electroporation and subsequently transformed into *A. thaliana ffc* background by floral dip (Clough and Bent, 1998). The homozygous plant lines were selected as described in Lehmann et al., 2017 and identified via genotyping PCR (Supplementary Figure S1A). All PCR products were checked by Sanger sequencing. Used primers are listed in Supplementary Table 1. The transgene expression was additionally verified by western blot (Supplementary Figure S1B).

### Plant material, growth conditions and phenotypic analyses

*Arabidopsis thaliana* wild type (Columbia-0, Col-0), *ffc* (*ffc* 1-2, previously described in (Amin et al., 1999)), and generated complementation lines (cpSRP54/*ffc*, cpSRP54NG/*ffc*, cpSRP54ΔC/*ffc*, cpSRP54(Q185R)/*ffc*) were grown on soil under artificial light (Philips Master TL-D 58W/840 Reflex Eco: 8 h light, 120 µE/m^-2^s^-1^, 22°C 65% humidity, 16 h dark, 19,5°C, 65% humidity). Total chlorophyll content of 100 mg Arabidopsis leaves from 4-week-old plants was determined using 80% (v/v) acetone according to Porra et al., 1989. The diameter of rosette size and their weight was measured from 10-week-old plants.

### Measurement of the maximum quantum yield of PSII

The maximum quantum yield of PSII (Fv/Fm) was analyzed using the Dual-PAM-100 measuring system (Walz GmbH, Germany). The 8-week-old Arabidopsis plants were acclimated in the dark for 30 min prior measurement. Fv/Fm was determined either after normal light (120 µE/m^-2^s^-1^) or 2 h high light stress (1200 µE/m^-2^s^-1^).

### Arabidopsis chloroplast isolation and fractionation

Isolation and fractionation of intact chloroplasts was done as previously described (Hristou et al., 2019; Stolle et al., 2022). Lysis of the chloroplasts was carried out at 1 to 2 mg chlorophyll/ml in HM buffer (10 mM magnesium chloride, 50 mM HEPES pH 8.0) for 30 min on ice and thylakoids were separated from stroma by centrifugation (20000 g, 4°C, 10 min). For thylakoid samples these were finally washed in HM buffer containing 150 mM sodium chloride and then adjusted to the desired chlorophyll concentration.

### Blue-native-polyacrylamide gel electrophoresis

Thylakoids isolated from chloroplasts of *Arabidopsis thaliana* plants (Col-0, *ffc*, cpSRP54/*ffc*, cpSRP54NG/*ffc*, cpSRP54ΔC/*ffc*, cpSRP54(Q185R)/*ffc*) were solubilized with 1.5% (w/v) n-dodecyl-β-D-maltosid (DDM) at a chlorophyll concentration of 1 mg/ml for 20 min on ice. Following centrifugation (20000 g, 4°C, 10 min) the BN-PAGE samples were prepared using NativePAGE™ 5% G-250 sample additive (Invitrogen) at a final concentration of 0.25%. The multiprotein complexes were separated by a NativePAGE™ 4-16% blue-native-polyacrylamide gel electrophoresis (BN-PAGE) gradient according to the manufacturer’s instructions (Invitrogen) using total amounts of 10 µg chlorophyll/lane.

### Two-dimensional BN-PAGE/SDS-PAGE followed by immunoblot analysis

Multiprotein complexes of thylakoids from *Arabidopsis thaliana* plants (Col-0, *ffc*, cpSRP54ΔC/*ffc*) were separated by BN-PAGE as described previously with adjusted application quantity according to the chlorophyll determination. Gel strips of the first dimensional NativePAGE™ 4-16% BN-PAGE gradient gel were incubated in preparation buffer (63 mM Tris pH 6.8, 33% (v/v) glycerol, 6.6% (w/v) SDS, 6.6% (v/v) β-mercaptoethanol, 6 M Urea) for 1 h at 4°C. For two-dimensional (2D) SDS-PAGE, multiprotein complexes were subsequently fractionated by electrophoresis on denaturing SDS gels (15% acrylamide) supplemented with 6 M urea (Schägger and Jagow, 1987). 2D gels were either stained with Coomassie® Brilliant Blue R-250 or proteins were transferred onto PVDF membranes by western blot. The dye was removed from the membrane by incubation in methanol. The membranes were subjected to immunoblot analysis with antibodies against PsbA, PsbD, PsaA, PsaB, PsaF, PsaK, Lhcb1, Lhca1, Lhca2, Lhca3, Lhca4, and PetA.

### Distribution determination of PSI core proteins in PSI* and PSI

ImageJ was used to determine protein levels of PsaA and PsaB on immunoblot band intensities after 2D SDS-PAGE. The measured values were calculated in relation to PSI* and PSI combined as 100%. Data was visualized in GraphPad Prism (v9.2.0).

### Immunoblot analyses

Proteins separated by SDS-PAGE were blotted onto nitrocellulose or PVDF membranes. The immunological detection was performed using specific antibodies against PsbA (AS05 084), PsbD (AS06 146), PsbO (AS06 142-33), PsaA (AS06 172), PsaB (AS10 695), PsaF (AS06 104), PsaK (AS04 049), Lhcb1 (AS01 004), Lhcb4 (AS04 045), Lhca1 (AS01 005), Lhca2 (AS01 006), Lhca3 (AS01 007), Lhca4 (AS01 008), PetA (AS08 306), uL4 (AS22 4787) obtained from Agrisera, His-tag (Penta-His HRP Conjugate, ID 34460, Qiagen), Actin (A0480, Merck), cpSRP54M (Walter et al., 2015), cpSRP43 (Klimyuk et al., 1999).

### Total protein extracts of Arabidopsis leaves

Total protein extracts of 100 mg Arabidopsis leaves from 4-week-old plants (Col-0, *ffc*, cpSRP54/*ffc*, cpSRP54NG/*ffc*, cpSRP54ΔC/*ffc*, cpSRP54(Q185R)/*ffc*) were prepared using TRIzol reagent (Life Technologies). Leaves were snap-frozen in liquid nitrogen and homogenized in 1 ml TRIzol reagent using a pistil. Samples were mixed with 200 µl chloroform and incubated for 10 min at room temperature (RT) followed by centrifugation (12000 g, RT, 5 min). The pellets were resuspended in 300 μl 100% (v/v) ethanol, incubated for 3 min at RT and again centrifuged (2000 g, 4°C, 15 min). The resulting pellet was discarded, and the supernatant mixed with 1 ml isopropanol. After rotating incubation for 30 min at RT and centrifugation (12000 g, 4°C, 10 min), the protein pellets were washed three times with 0.3 M guanidine hydrochloride in 95% (v/v) ethanol. A final washing step was carried out using 2 ml 100% (v/v) ethanol. Proteins were pelletized by centrifugation (7600 g, 4°C, 5 min) and subsequently dried at 75°C and 750 rpm. Protein pellets were resolved overnight in 100 µl 1% (w/v) SDS ultrapure followed by centrifugation (10000 g, RT, 10 min) to get rid of insoluble compounds. The total protein extracts were mixed with sample buffer and subsequently applied to SDS-PAGE and Western Blot analyses. Samples were adjusted to actin content for further immunoblot analyses. The protein levels of PsbA, PsbO, PsaA, PsaB, Lhcb1, Lhca2 and Actin were quantified based on immunoblot band intensities using ImageJ in relation to 100% wild type total protein extract.

### Statistical analyses

Statistical analyses and data visualization were performed in GraphPad Prism (v9.2.0). To determine statistically significant differences between means, one– or two-way analysis of variance (ANOVA) was followed by Sidak’s or Dunnett’s multiple comparisons test.

### Isothermal titration calorimetry (ITC)

The thermodynamic values of recombinant cpSRP54 and cpSRP54(Q185R) binding to GDP and GTP were measured using a MicroCal ITC200 instrument (Malvern Panalytical). Protein and ligand concentrations were determined using a NanoDrop One spectrophotometer (Thermo Fisher Scientific). 25 µM of protein and 1 mM of ligand, diluted in SEC buffer (20 mM HEPES, pH 7.5, 200 mM NaCl, 20 mM MgCl_2_, and 20 mM KCl), were loaded in the cell and syringe, respectively. All measurements were done at 25°C at a stirring rate of 750 rpm applying 1 injection of 0.4 µl and 12 more of 3 µl with a spacing of 150 s. Raw data was analyzed in the MicroCal PEAQ-ITC Analysis Software v1.41 (Malvern Panalytical) applying the One Set of Sites model. The resulting thermodynamic parameters can be found in Supplementary Table 2.

### Determination of GTPase activity

The GTPase activity of recombinant cpSRP54 and cpSRP54(Q185R) in the presence of cpFtsY-NG was assayed in a buffer containing 25 mM HEPES (pH 7.5), 10 mM Mg(OAc)_2_, 300 mM K(OAc), 1 mM DTT, and 2.5% (v/v) glycerol. Enzymatic reactions containing 10 µM of each protein and 1 mM of GTP were incubated at 37°C for up to 60 min. Samples taken at each corresponding time point were quenched by adding chloroform, vortexing and subjecting them to 95°C for 15 s after which they were snap-frozen in liquid nitrogen. Thawed samples were centrifuged at 13000 g for 5 min at 4°C and the resulting aqueous phase was then analyzed by high-performance liquid chromatography (HPLC) in an Agilent 1260 Infinity system (Agilent) using a Metrosep A Supp 5 – 150/4.0 column (Metrohm). A flow rate of 0.6 ml/min in a buffer containing 90 mM (NH_4_)_2_CO_3_ (pH 9.25) was used and a wavelength of 260 nm was selected for nucleotide detection. GDP and GTP samples (Jena Bioscience) were used as standards to identify the retention time of these two nucleotides within the experimental samples. GTP consumption was determined by comparing the peak area corresponding to GTP at time = 0 to the following time points.

### Protein Expression and purification of recombinant proteins

Gene fragments encoding residues 65-366 of *A. thaliana* cpFtsY and residues 79-564 of *A. thaliana* cpSRP54 were cloned via GoldenGate (NEB) into a pET24d derivative (AL324, Novagen) modified for modular cloning for overproduction of C-terminal His6-tagged proteins (Kohm et al., 2023). CpFtsY was cloned via BsaI and cpSRP54 via Esp3I restriction sites resulting in AL324_cpFtsY-65 and AL324_cpSRP54 constructs. The overexpression constructs encoding mature cpFtsY and mature cpSRP54 described previously were used as templates (Bals et al., 2010). The construct cpSRP54(Q185R) was generated using the QuikChange Lightning site-directed mutagenesis kit (Agilent Technologies) with pETDuet1™-cpSRP54 (Bals et al., 2010) serving as template. Primers are listed in Supplementary Table 1. The correct sequence of the constructs was verified by sequencing. His-tagged recombinant proteins were overexpressed in *Escherichia coli* strain Rosetta™ (DE3) and purified via an Äkta chromatography system using HisTrap columns (GE Healthcare, Waukesha, WI, USA). Washing (20 mM HEPES pH 8.0, 250 mM NaCl, 20 mM KCl, 40 mM imidazole) and elution (20 mM HEPES pH 8.0, 250 mM NaCl, 20 mM KCl, 250 mM imidazole) buffers did not contain magnesium chloride. Elution fractions were collected and supplemented with 30 mM EDTA for 20 min at RT. After concentration using centrifugal filters (Amicon Ultra, Witten, Germany), the elution was subjected to a size exclusion chromatography (column XK16 S200, GE Healthcare, Waukesha, WI, USA) using a magnesium chloride containing running buffer (20 mM HEPES pH 7.5, 200 mM NaCl, 20 mM MgCl_2_, 20 mM KCl). The elution fractions of the corresponding protein peak were pooled and concentrated as before, and the purified proteins were used for subsequent experiments.

## Results

### Full complementation of the developmental defect of the ffc mutant requires expression of wild-type cpSRP54

To explore the function of the cpSRP54 protein *in vivo*, we generated four *A. thaliana* complementation lines of the cpSRP54-knockout mutant *ffc1-2* (in the following referred to as *ffc*) (Amin et al., 1999) using constructs encoding His-tagged cpSRP54 variants under the control of the 35S promoter. The complementation of the *ffc* mutant, which is reduced in growth and shows a chlorotic phenotype, particularly at a young developmental stage, was analyzed by the expression of full-length cpSRP54 (cpSRP54/*ffc*), protein variants lacking the complete C-terminal half including the methionine-rich M-domain (cpSRP54NG/*ffc*) or its C-terminal tail region (cpSRP54ΔC/*ffc*), and a point mutation variant, where a glutamine (Q) of the phosphate (P)-binding loop of the GTPase motif I (G1) in the G-domain was replaced by an arginine (R) (cpSRP54(Q185R)/*ffc*) (Figure 1A). We selected this point mutation because previous studies demonstrated a detrimental effect of a mutation at this position on GTP hydrolysis in bacterial SRP GTPases (Samuelsson et al., 1995; Egea et al., 2004; Shan et al., 2004). After *Agrobacterium tumefaciens*-mediated transformation of the *ffc* plants, we screened the progeny of each transformation event, and lines with clearly measurable expression levels of the cpSRP54 variants were used for further analyses (Figure 1B, Supplementary Figure S1A/B). Furthermore, we confirmed the presence of the cpSRP54-His variants in stromal extracts of the plant lines by immunoblot analyses and verified their expected migration behaviour by sucrose gradient centrifugation of the extracts (Supplementary Figure S2).

The cpSRP54/*ffc* lines showed full complementation of the *ffc* mutant phenotype and exhibited similar rosette sizes and weights and even a slightly increased chlorophyll content compared to wild type plants (Figure 1B, Figure 2A-C). In contrast, the cpSRP54NG/*ffc* as well as the cpSRP54(Q185R)/*ffc* lines resembled the *ffc* mutant showing significantly smaller and lighter plant rosettes than those of wild type (Figure 1B, Figure 2A/B). Comparable to the *ffc* mutant, the chlorophyll content was reduced to 56% and 54% of wild type level in cpSRP54NG/*ffc* and cpSRP54(Q185R)/*ffc,* respectively (Figure 1B, Figure 2C). The cpSRP54ΔC/*ffc* plants, however, displayed a partial phenotypic complementation of the *ffc* mutant with more developed rosettes (Figure 1B, Figure 2A/B) and a total chlorophyll content of 82% of wild type (Figure 2C). Despite this, the overall phenotypic analysis of cpSRP54ΔC/*ffc* revealed significant differences compared to the wild type plants (Figure 2A-C).

**Figure 2:**
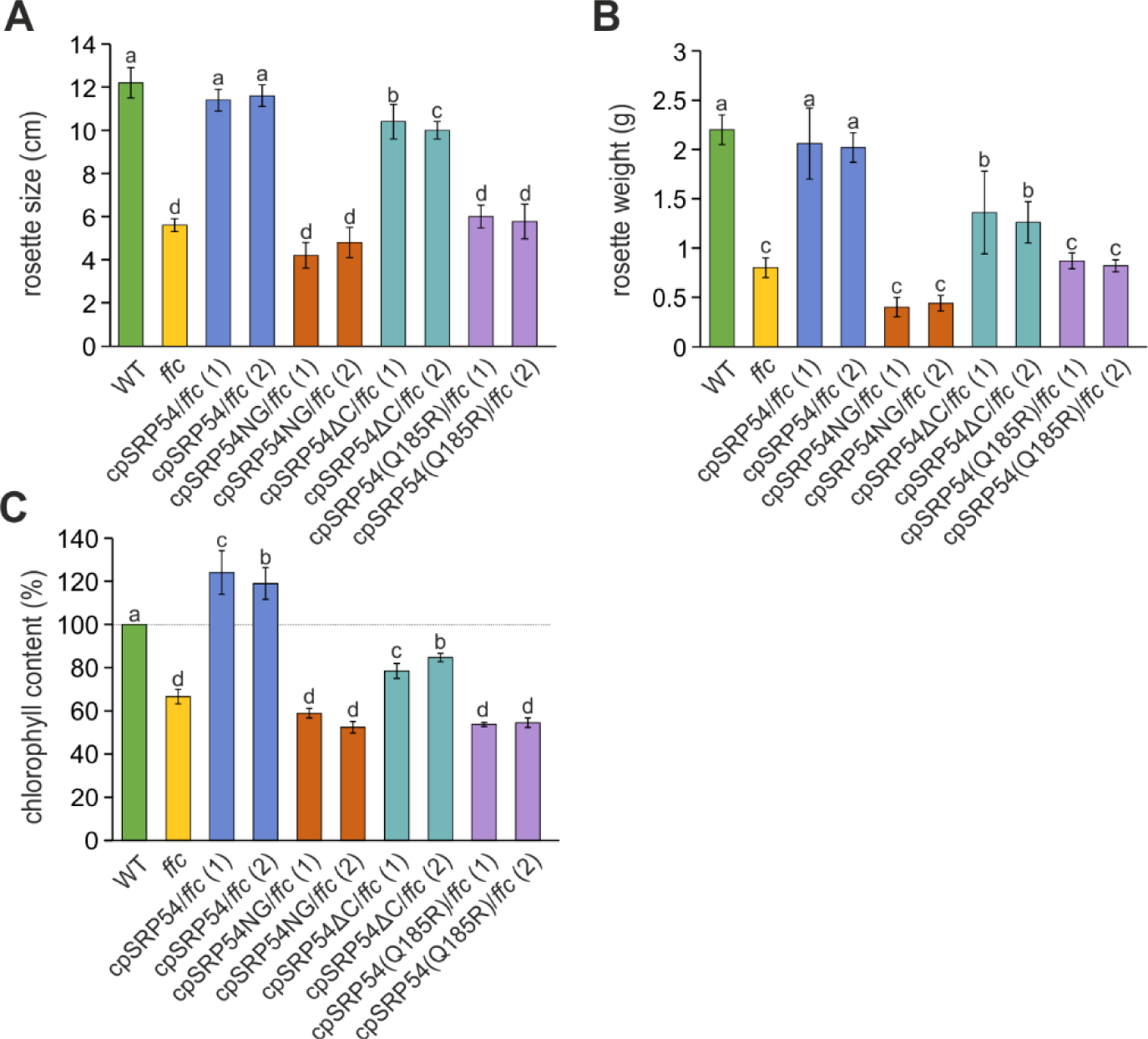
*A. thaliana ffc*-complementation lines display altered phenotypes. The rosette size. (**A**) and rosette weight (**B**) was measured from 10-week-old plants. (**C**) The chlorophyll content was determined from 4-week-old *A. thaliana* plants in relation to wild type (WT, dotted line (100%)). In each analysis the average values and corresponding SDs were calculated from three to eight independent measurements. Different letters indicate statistically significant differences (p < 0.05) between means determined by one-way ANOVA followed by Dunnett’s multiple comparisons test.

### Impaired PSII activity of the ffc mutant is rescued by cpSRP54 lacking its C-terminal tail region, but not by cpSRP54 variants with a functionally impaired G-domain or lacking the M-domain

Since the *ffc* mutant shows a defect in PSII repair (Walter et al., 2015b) we set out to determine the functionality of PSII in the complementation lines compared to wild type and *ffc.* Therefore, the maximum quantum efficiency of PSII in the dark-adapted state (Fv/Fm) was measured after the plants were exposed to normal (NL) and high light (HL) conditions. As expected, the cpSRP54/*ffc* plants showed Fv/Fm values similar to wild type under both light conditions (Figure 3A). In contrast to wild type and the fully complemented lines, *ffc* as well as cpSRP54NG/*ffc* and cpSRP54(Q185R)/*ffc* showed significant differences in the Fv/Fm values between the NL and HL measurements indicating that high light treatment leads to a strong impairment of PSII activity in these plants (Figure 3A). Here, the response of the cpSRP54NG/*ffc* and cpSRP54(Q185R)/*ffc* plants to high light was even more pronounced than that of *ffc* (Figure 3B). Interestingly, expression of cpSRP54ΔC in the *ffc* mutant complemented the impaired PSII activity of the *ffc* plants and even under high light stress the cpSRP54ΔC/*ffc* plants showed no significant difference in PSII activity compared to wild type (Figure 3A/B). Together, the measurements suggest that the C-terminal tail region of cpSRP54 is not essential for biogenesis and maintenance of PSII, while the M-domain and a functional G-domain play important roles in these processes.

**Figure 3:**
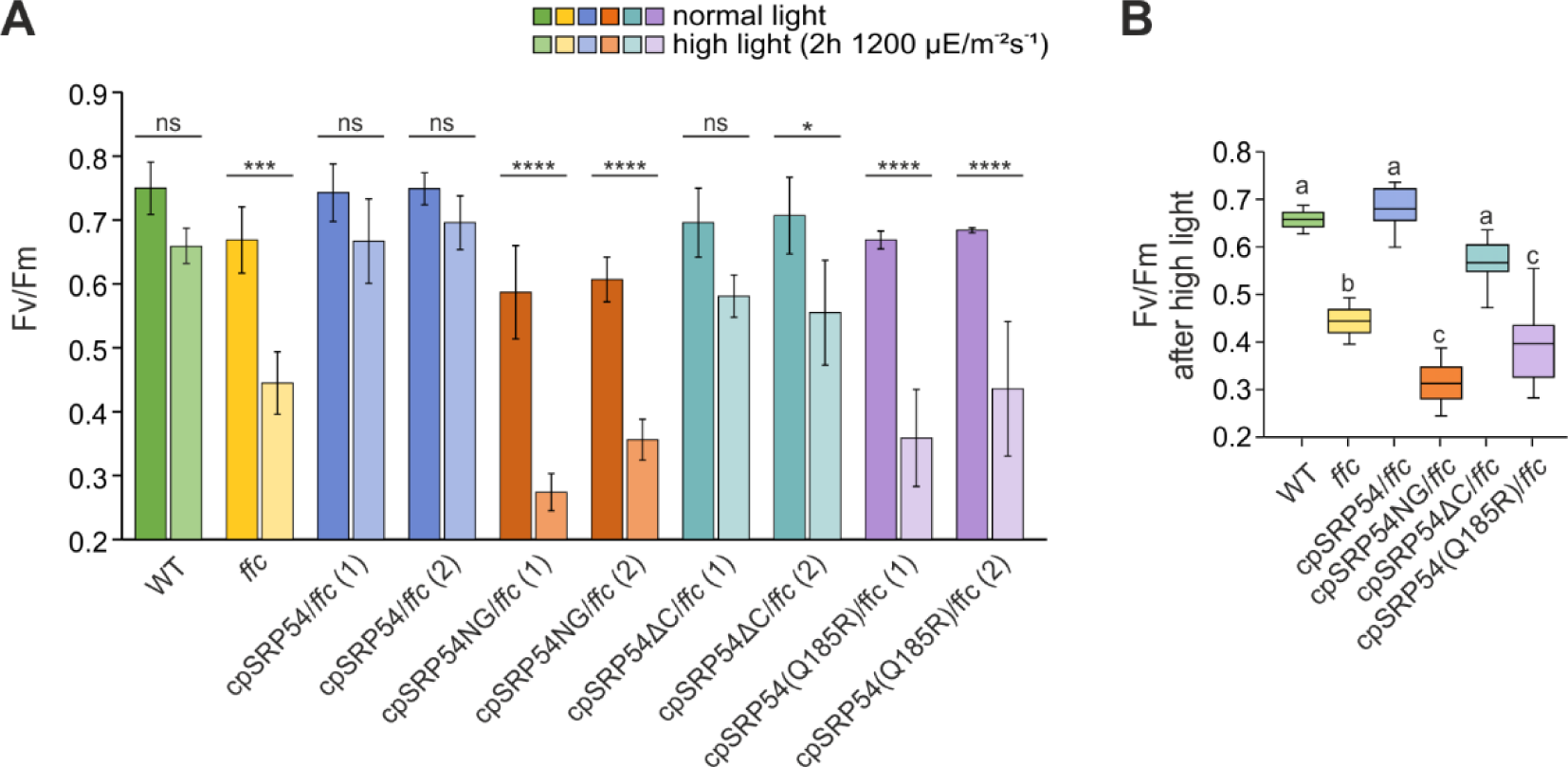
Measurement of the photosynthetic performance in *A. thaliana* wild type, *ffc* and the *ffc*-complementation lines. The maximum quantum yield of PSII (Fv/Fm) of 8-week-old wild type (WT), *ffc*, and *ffc*-complementation plants was measured under constant normal light conditions and after 2h high light stress. The Fv/Fm values were calculated from three to five independent measurements. (**A**) The means of normal and high light conditions were compared using two-way ANOVA followed by Sidak’s multiple comparisons test. Statistically significant differences are marked by asterisks (*p <0.05, **p < 0.01, ***p < 0.001, ****p < 0.0001, ns = no significance). (**B**) The means of the measured Fv/Fm after high light treatment were analyzed using one-way ANOVA followed by Dunnett’s multiple comparisons test. Different letters indicate statistically significant differences (p < 0.05) between means.

### The cpSRP54 C-terminal region as well as functional G– and M-domains are required for the posttranslational cpSRP transport of LHCPs

To determine the protein levels of PSI and PSII components in the thylakoid membrane of the complementation lines, *ffc* and wild type, we performed an immunoblotting analysis using total protein extracts. This and all further experiments were carried out using one of the two complementation lines investigated so far. Here, we focused on the plastid-encoded PSI and PSII core subunits (PsaA, PsaB and PsbA) and nuclear-encoded LHCPs (Lhca2 and Lhcb1), which are transported to the thylakoid membrane using the co– or posttranslational cpSRP-dependent mechanism, respectively. The cytoplasmic protein Actin and the lumenal PSII subunit PsbO, which is transported via the cpSec1-dependent protein transport pathway, served as controls (Schünemann et al., 1999). Quantification of the detected protein levels revealed that the observed protein levels in *ffc* (Figure 4A) are in accordance with previous results (Amin et al., 1999; Rutschow et al., 2008; Hristou et al., 2019), showing levels between 45% and 52% for PsaA, PsaB and PsbA as well as 52% for Lhcb1 and 71% for Lhca2 compared to wild type (Figure 4B/C).

**Figure 4:**
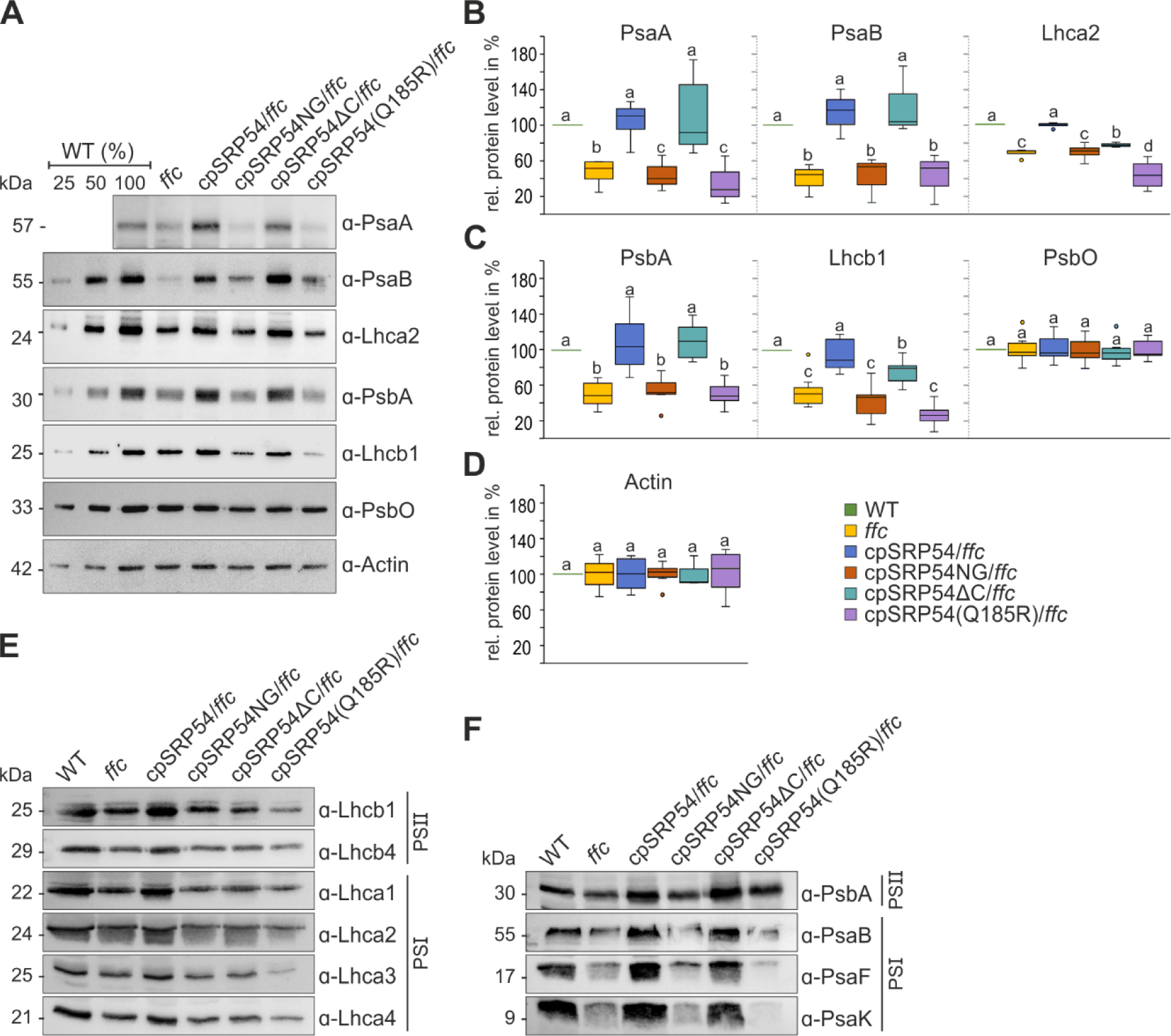
Immunoblot analysis of PSI and PSII subunits and LHCPs in total protein extracts and thylakoid membranes of *A. thaliana* wild type, *ffc* and the *ffc*– complementation lines. (**A**) Total protein extracts from fresh leaf material of 4– to 5-week-old wild type (WT), *ffc* and *ffc*-complementation plants were separated by SDS-PAGE and immunoblotted with the indicated antibodies. The WT extracts correspond to 25, 50, and 100% of total protein. The protein levels of PSI (**B**) and PSII (**C**) proteins, and the cytoplasmic protein Actin (**D**) were quantified by immunoblot band intensities using ImageJ in relation to wild type (100%). Means were calculated from three to nine independent replicates. Different letters indicate statistically significant differences (p < 0.05) between means determined by one-way ANOVA followed by Dunnett’s multiple comparisons test. (**E, F**) Isolated thylakoids from 4– to 5-week-old WT and the transgenic lines were solubilized in DDM and analyzed by SDS-PAGE followed by immunoblot with the indicated antibodies against LHCPs (E) and (F) photosystem subunits. The samples were separated based on the measured relative chlorophyll content of the plant lines (WT: 100%, *ffc*: 67%, cpSRP54/*ffc*: 124%, cpSRP54NG/*ffc*: 59%, cpSRP54ΔC/*ffc*: 79%, cpSRP54(Q185R)/*ffc*: 54%).

The cpSRP54NG/*ffc* and cpSRP54(Q185R)/*ffc* lines show no complementation of the PSI and PSII core subunits and the LHC proteins at all (Figure 4B/C). A full recovery of the wild type protein level of all photosynthetic proteins was exclusively observed in the cpSRP54/*ffc* plants (Figure 4A) exhibiting protein amounts of 104-117% of PsaA, PsaB and PsbA, and 90-100% of Lhcb1 and Lhca2 compared to wild type (Figure 4B/C). Interestingly, the cpSRP54ΔC/*ffc* also shows pronounced complementation of the PSI and PSII reaction center proteins (92% PsaA, 104% PsaB and 109% PsbA of wild type level) but much less pronounced complementation in the level of Lhcb1 (75% of wild type level) and almost no complementation in the level of Lhca2 (77% of wild type level) (Figure 4A-C). Since the use of total leaf protein extracts may involve the detection of proteins that are not integrated into the thylakoid membrane but accumulate in soluble fractions, immunoblotting experiments were performed using solubilized thylakoid membranes with chlorophyll concentrations based on the previously measured relative chlorophyll content of the plant lines (Figure 2C). The analyzed thylakoid membranes show clear reductions in a variety of PSI and PSII LHCPs (PSI: Lhca1-4, PSII: Lhcb1, Lhcb4) in *ffc* and all complementation lines except cpSRP54/*ffc* (Figure 4E). Moreover, while *ffc*, cpSRP54NG/*ffc* and cpSRP54(Q185R)/*ffc* showed a reduction of the PSI and PSII reaction center subunits PsaB and PsbA, these subunits were markedly upregulated in cpSRP54/*ffc* and cpSRP54ΔC/*ffc*, displaying protein amounts comparable to wild type (Figure 4F).

Taken together, the results demonstrate that the C-terminus of cpSRP54 plays an important role for the posttranslational transport of LHCPs but is not required for cotranslational transport of the PSI and PSII reaction center proteins *in vivo*. Furthermore, our data indicate that the M-domain and the GTPase activity of the G-domain are essential for the biogenesis of the PSI and PSII cores as well as for the LHCI and LHCII antenna systems.

### The loss of the cpSRP54 C-terminus promotes the accumulation of immature PSI

To obtain insight into the assembly of thylakoid membrane complexes in the cpSRP54 complementation lines, isolated thylakoids were treated with n-dodecyl-β-D-maltoside (DDM), and the solubilized complexes subsequently fractionated by blue-native (BN)-PAGE. To analyze the relative abundances of the complexes in the complementation lines, samples were loaded according to the proportional chlorophyll concentrations of the plant lines with wild type serving as 100% control (Figure 5). The complex distribution of wild type and cpSRP54/*ffc* showed similar patterns with four distinct PSII supercomplexes (SC), fully assembled PSI and PSII dimers, monomeric PSII with the directly above running Cyt*b*_6_*/f* complex, the CP43-free PSII monomer, the reaction center-like complex as well as multimeric, trimeric, and monomeric light-harvesting antenna complexes II (LHCII) at the expected sizes (Figure 5). The same pattern of complexes, although in reduced amounts, was observed in *ffc* as well as in the complementation lines cpSRP54NG/*ffc*, cpSRP54(Q185R)/*ffc*, and cpSRP54ΔC/*ffc*. Strikingly, however, cpSRP54ΔC/*ffc* showed a clear accumulation of a complex (indicated as (III) in Figure 5) with a running behaviour similar to the previously described PSI* complex, representing an intermediate assembly state during PSI biogenesis in *N. tabacum* (Wittenberg et al., 2017). To analyze the subunit composition of the accumulated complex in cpSRP54ΔC/*ffc* and to identify further potential differences in the photosynthetic complexes of wild type and cpSRP54ΔC/*ffc*, BN-PAGE slices were subsequently subjected to SDS-PAGE analysis (2-dimensional (2D) BN/SDS-PAGE) followed by coomassie staining or immunoblot analyses (Figure 6A/B). As the plastid-encoded Cyt*b*_6_*/f* complex subunit PetA is targeted independent from cpSRP54 by a pathway involving cpSecA1, it served as experimental control (Röhl and van Wijk, 2001). In cpSRP54ΔC/*ffc* the PSII core subunits (PsbA and PsbD) were reduced in the PSII SC, while the levels of PSII dimeric and monomeric complexes as well as the CP43-free PSII assembly intermediate are similar to wild type. We also observed a reduction of Lhcb1 in the PSII SC in cpSRP54ΔC/*ffc*. Furthermore, the level of Lhcb1 present as LHCII trimers was clearly reduced in this mutant. These findings are consistent with the data described above and corroborate that the cpSRP54ΔC/*ffc* line is impaired in the biogenesis of LHCII, while the biogenesis of the PSII core is unaffected.

**Figure 5:**
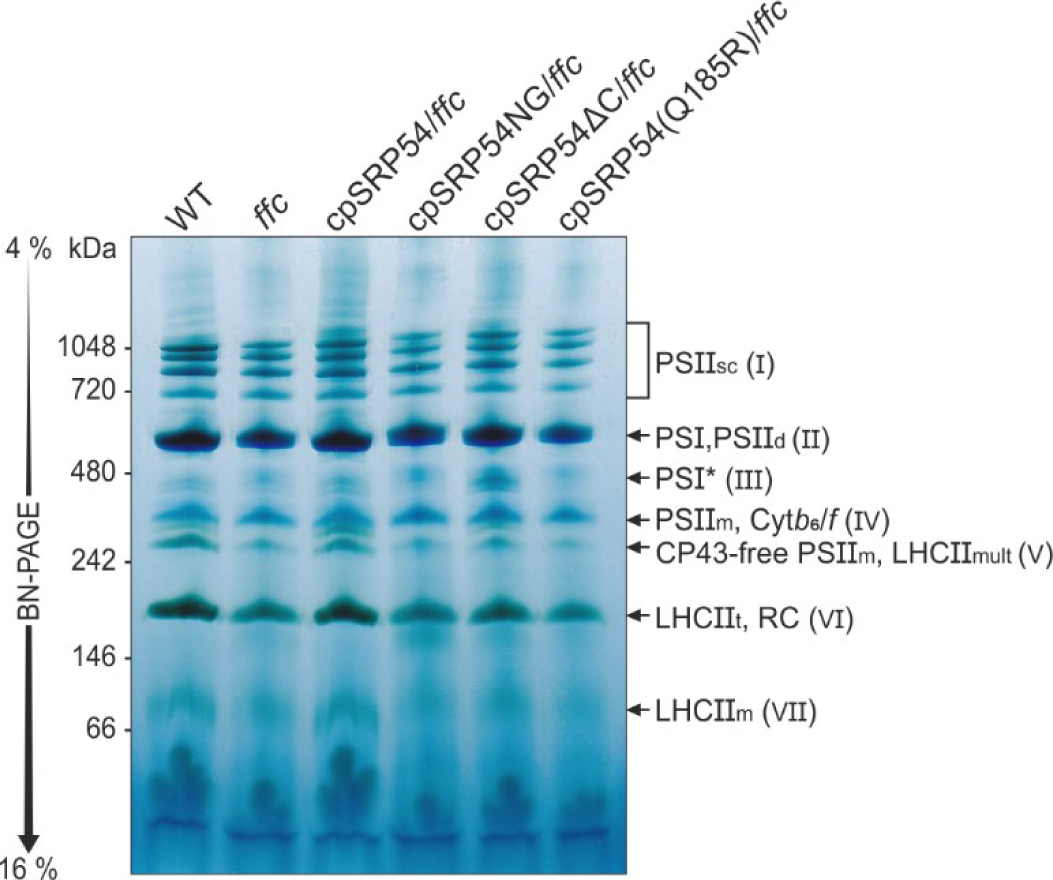
BN-PAGE analysis of thylakoid membrane multiprotein complexes of *A. thaliana* wild type and the transgenic lines. Isolated thylakoids from 4– to 5-week-old wild type (WT), *ffc* and *ffc*-complementation plants were solubilized in DDM and separated by BN-PAGE based on their measured relative chlorophyll content (WT: 100%, *ffc*: 67%, cpSRP54/*ffc*: 124%, cpSRP54NG/*ffc*: 59%, cpSRP54ΔC/*ffc*: 79%, cpSRP54(Q185R)/*ffc*: 54%). The identification of detected bands was accomplished in accordance with published BN-PAGE profiles of Arabidopsis thylakoids (Granvogl et al., 2006; Armbruster et al., 2010; Wittenberg et al., 2017; Che et al., 2022). Photosystem I and II (PSI and II), PSII supercomplexes (PSII_sc_, I), PSII dimers (PSII_d_, II), PSI assembly intermediate (PSI*, III), monomeric PSII (PSII_m_, IV) and cytochrome *b*_6_*/f* complex (Cyt*b*_6_*/f*, IV), CP43-free PSII monomers (CP43-free PSII_m_, V) and multimeric light harvesting antenna complex II (LHCII_mult_, V), trimeric LHCII (LHCII_t_, VI), reaction center-like complex (RC, VI), monomeric (LHCII_m_, VII).

**Figure 6:**
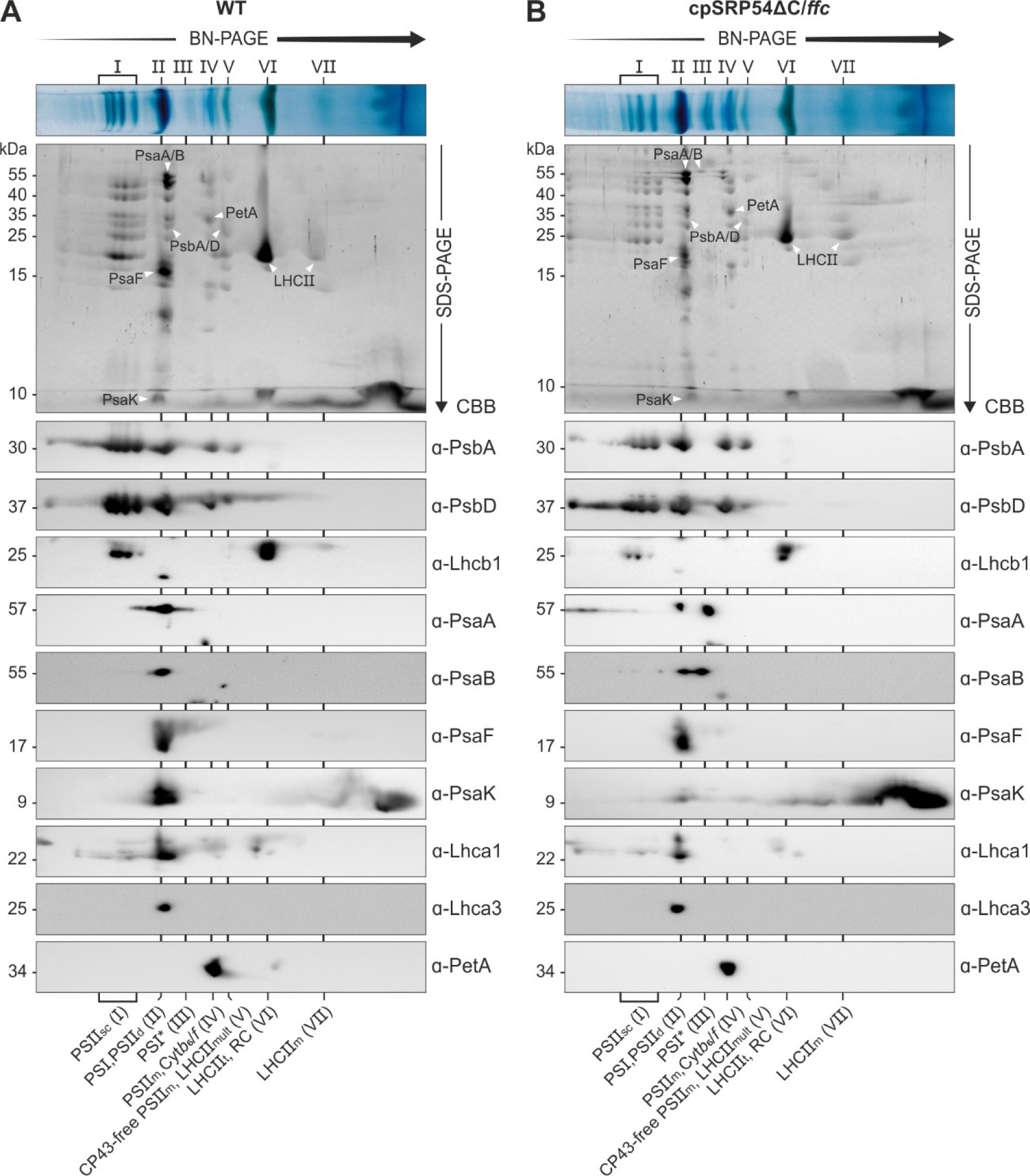
Two dimensional BN-PAGE/SDS-PAGE analysis of thylakoid membrane multiprotein complexes in *A. thaliana* wild type and the cpSRP54ΔC/*ffc*-complementation line. Thylakoids of 4– to 5-week-old plants were solubilized in DDM and separated by BN-PAGE based on their measured relative chlorophyll content ((**A**) wild type (WT) 10 µg chlorophyll/lane, (**B**) cpSRP54CΔC/*ffc* 7.9 µg chlorophyll/lane). Detected bands were identified in accordance with published BN-PAGE profiles of Arabidopsis thylakoids (see legend to Figure 6). The protein complex subunit composition was further determined by two-dimensional (2D) BN-PAGE/SDS-PAGE analysis followed by Coomassie blue staining (CBB) or western blot with subsequent immunodetection using the indicated antibodies.

Notably, the PSI core subunits (PsaA and PsaB) displayed two clearly separated antibody signals in cpSRP54ΔC/*ffc*, while only one signal was primarily detected in wild type (Figure 6B). The signals could be assigned to the fully assembled PSI and the accumulated complex in cpSRP54ΔC/*ffc*. (Figure 5).

Next, we wanted to test whether the subunits PsaF and PsaK, which were described to be absent or to occur only in low abundances in the PSI* complex of *N. tabacum* (Wittenberg et al., 2017), are present in the PSI assembly intermediate. Both PSI subunits were reported to incorporate into the thylakoid membrane following the assembly of the PSI reaction center (Ozawa et al., 2010; Wittenberg et al., 2017; Zhang et al., 2024). Immunodetection of PsaF and PsaK using solubilized thylakoid membranes showed no difference in the total amount of PsaF and PsaK between wild type and cpSRP54ΔC/*ffc* (Figure 4 F). The 2D BN/SDS-PAGE immunoblot signals revealed that PsaF as well as PsaK are present in the fully assembled PSI but not in the PSI assembly intermediate. Remarkably, the amount of PsaK detected in the fully assembled PSI was drastically reduced in cpSRP54ΔC/*ffc* while an accumulation as non-complexed protein was observed (Figure 6B).

We also tested antibodies against LHCPs (Lhca1 and Lhca3) to determine whether the *A. thaliana* PSI assembly intermediate is associated with antenna proteins, which are absent in the PSI* complex of *N. tabacum* (Wittenberg et al., 2017). As shown in Figure 6B, no comigration of Lhca1 and Lhca3 with the PSI assembly intermediate was detected (Figure 6B), while fully assembled PSI is associated with Lhcas to some extent. Notably, we were also able to detect small amounts of the PSI assembly intermediate in the *ffc* mutant (Supplementary Figure S3). The observed antibody signals in the immunoblots after 2D BN/SDS-PAGE are consistent with the results of the analysis described above. Furthermore, immunoblots using antibodies against Lhca2 and Lhca4 clearly confirm that the *A. thaliana* PSI assembly intermediate is not associated with LHCPs (Supplementary Figure S3). In conclusion, the PSI assembly intermediate identified in *A. thaliana* shares properties with the PSI* complex from *N. tabacum* and is henceforth referred to as PSI*.

To analyze the relative abundance of the PSI and PSI* complexes in wild type, *ffc* and cpSRP54ΔC/*ffc*, the signals of the PSI core subunits in the immunoblots were quantified. While in wild type the vast majority was assigned to the fully assembled PSI (PSI: 94% PsaA, 95% PsaB), the distribution in *ffc* was altered (PSI: 73% PsaA, 81% PsaB; PSI*: 27% PsaA, 19% PsaB) and in cpSRP54ΔC/*ffc* even markedly shifted towards the PSI* complex (PSI: 40% PsaA, 47% PsaB; PSI*: 60% PsaA, 53% PsaB) (Figure 7).

**Figure 7:**
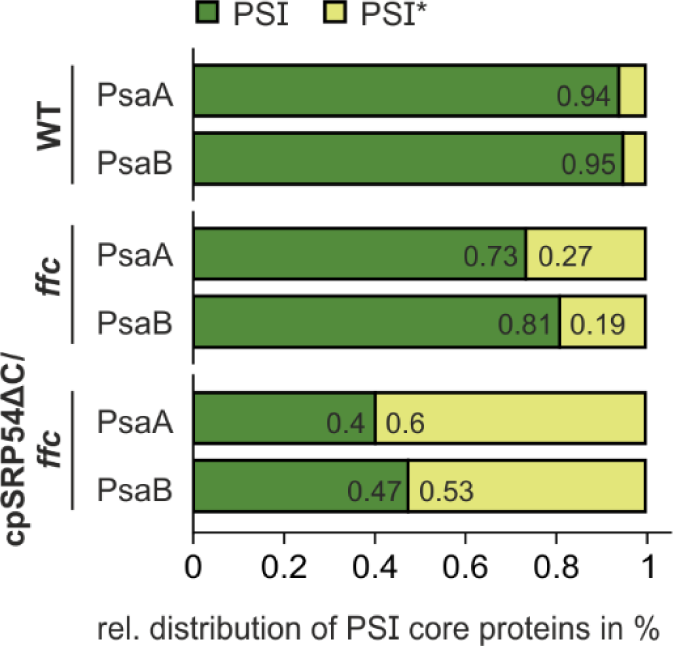
Distribution of the PSI core protein subunits. The protein levels of the PsaA and PsaB in the PSI* assembly intermediate and the fully assembled PSI were determined by 2D BN-PAGE/SDS-PAGE immunoblot band intensities using ImageJ (n = 2-3). The total immunoblot signal calculated from both, PSI* and PSI, served as 100%.

Taken together, our data show that the impaired biogenesis of LHC proteins combined with the ongoing cotranslational formation of the PSI core, as seen in the cpSRP54ΔC/*ffc* line, leads to a strong accumulation of PSI*. Furthermore, our data suggest that the incorporation of PsaK in PSI is dependent on the addition of the LHC proteins during PSI assembly.

### Point mutation in the P-loop of cpSRP54 decelerates its GTPase activity

As our data showed that the cpSRP54(Q185R)/*ffc* plants exhibit a drastic impairment in plant development and significantly reduced levels of photosynthetic proteins and high-molecular protein complexes, the behaviour of cpSRP54(Q185R) was further investigated *in vitro*. Therefore, we quantitatively analyzed the interaction of recombinant cpSRP54-His and cpSRP54(Q185R)-His with either GTP or GDP using isothermal titration calorimetry (ITC). The measurements using cpSRP54-His resulted in an average dissociation constant of 55.2 µM (± 9.74) with GTP (Figure 8A), and 30.8 µM (± 5.78) with GDP (Figure 8B). For the titration of cpSRP54(Q185R)-His an average dissociation constant of 17.5 µM (± 2.86) with GTP (Figure 8C), and 30.2 µM (± 9.12) with GDP (Figure 8D) was determined. Despite the affinity of GDP being the same for both proteins, the affinity of GTP for the point-mutated protein cpSRP54(Q185R)-His was ∼3-fold higher than for the wild type protein cpSRP54-His.

**Figure 8:**
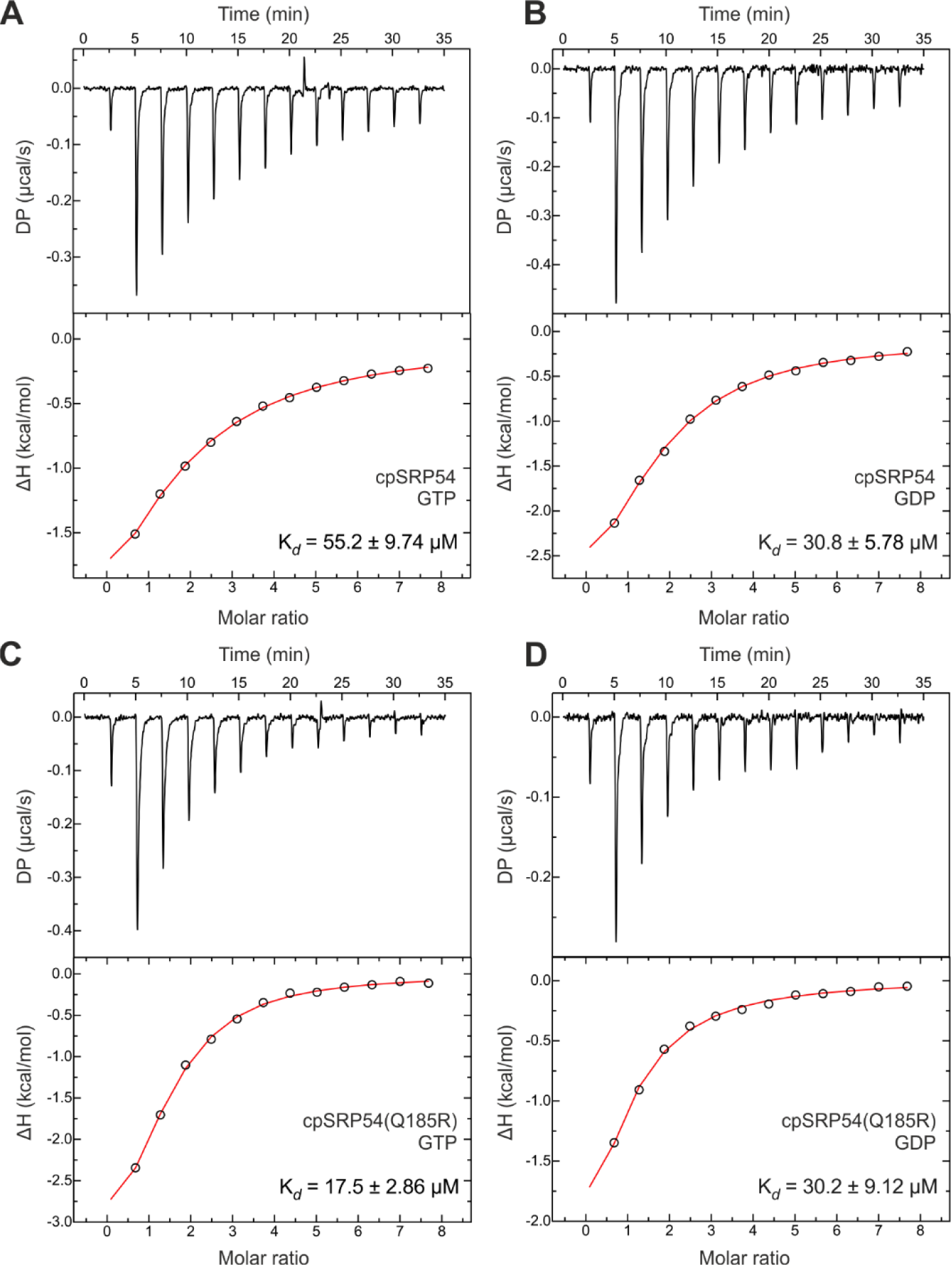
Isothermal titration calorimetry (ITC) measurement to determine binding affinities for the cpSRP54 protein and the point mutant cpSRP54(Q185R) with GTP and GDP. 1 mM of the ligand (GTP or GDP) was titrated with 25 µM of cpSRP54-His or cpSRP54(Q185R)-His. The resulting changes in heating power were recorded and the enthalpy changes were plotted versus the molar ratio of cpSRP54 or cpSRP54(Q185R) and the corresponding nucleotides. The titration isotherms resulted in K_d_’s as indicated in (**A**) cpSRP54/GTP, (**B**) cpSRP54/GDP, (**C**) cpSRP54(Q185R)/GTP, and (**D**) cpSRP54(Q185R)/GDP.

Moreover, we determined the GTPase activity of wild type and the point mutated proteins using high-performance liquid chromatography (HPLC). As cpSRP54 and its receptor protein cpFtsY were described to stimulate their mutual GTPase activity through interaction of their homologous G-domains (Jaru-Ampornpan et al., 2009; Wild et al., 2016), the experiments were performed in presence of cpFtsYNG-His. The GTP consumption was determined at different time points and set in relation to the initial GTP amount. After 5, 10, and 20 minutes the cpSRP54 containing samples exhibited a GTP consumption of 67%, 83%, and 93%, respectively. After 60 minutes the GTP was hydrolyzed to almost 100% (Figure 9). In comparison, GTP consumption was greatly reduced in the cpSRP54(Q185R) containing samples. This was most prominent after relatively short incubation times of 5, 10, and 20 minutes, which resulted in the hydrolysis of 30%, 44%, and 65% GTP (Figure 9). Overall, our data indicate that the higher affinity of cpSRP54(Q185R) to GTP traps the protein more persistently in its GTP-bound state, which results in a reduced GTP hydrolysis rate.

**Figure 9:**
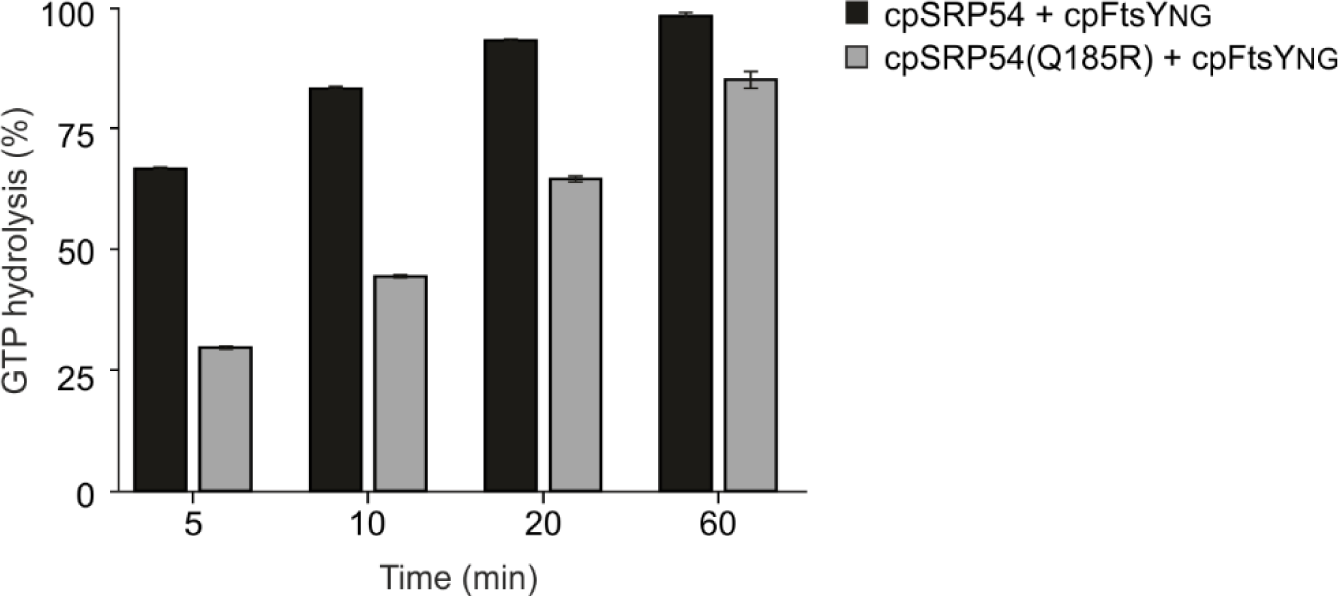
GTPase activity of cpSRP54 and cpSRP54(Q185R). The GTPase activity of cpSRP54-His and cpSRP54(Q185R)-His was measured by high-performance liquid chromatography (HPLC) in presence of cpFtsYNG-His using 10 µM of each protein and 1 mM of GTP (n = 3). The GTP hydrolysis was measured at different time points up to 60 min and the consumption was determined in relation to the 0 min time point.

To exclude the possibility that the point-mutation causes misfolding of the protein, a size exclusion chromatography was performed that demonstrated the same running behaviour for cpSRP54-His and cpSRP54(Q185R)-His (Supplementary Figure S4).

## Discussion

The role of the cpSRP54 protein within the SRP-dependent protein transport of higher plant chloroplasts has been characterized in several studies in the last decades. However, most studies were restricted to *in vitro* experiments and questions regarding the precise role of cpSRP54 *in vivo* remained open. Our study revealed a deeper insight into the functional role of the individual domains of cpSRP54 *in vivo.* The domains are similarly organized as in the homologous SRP54 proteins of eukaryotes and prokaryotes (Ffh), which mediate the cotranslational protein transport to the endoplasmic reticulum and the plasma membrane, respectively (Franklin and Hoffman, 1993; Grudnik et al., 2009; Akopian et al., 2013). However, cpSRP54 differs from its cytosolic homologues by a land plant-specific C-terminal tail region harboring the positively charged cpSRP43-binding motif (ARRKR in *A. thaliana*) (Funke et al., 2005). This motif is essential for the formation of the high affinity cpSRP54/cpSRP43 heterodimeric complex in *A. thaliana* (Ziehe et al., 2018). Furthermore, *in vitro* experiments to reconstitute transit complex formation with LHCP and insertion of LHCP into thylakoid membranes indicated that complex formation between the cpSRP subunits is essential for these processes (Schünemann et al., 1998; Tu et al., 1999; Yuan et al., 2002; Goforth et al., 2004; Funke et al., 2005). However, more recent *in vitro* studies suggested that cpSRP43 alone might be sufficient for LHCP targeting because it was demonstrated that cpSRP43 prevents LHCP aggregation by itself (Falk and Sinning, 2010; Jaru-Ampornpan et al., 2010) and is able to directly contact the Alb3 translocase in the thylakoid membrane (Bals et al., 2010; Falk et al., 2010; Lewis et al., 2010; Dünschede et al., 2011). A cpSRP43 mediated LHCP sorting was also indicated by the analysis of cpSRP-pathway Arabidopsis mutants. Notably, however, this pathway was only upregulated in the absence of cpSRP54 as well as its thylakoid receptor cpFtsY (Tzvetkova-Chevolleau et al., 2007). In this study, we show that a cpSRP54 variant, which lacks the C-terminal tail region is not able to rescue the defect in posttranslational LHCP transport in the *ffc* mutant. Therefore, our results are consistent with the *in vitro* LHCP insertion experiments and support the view, that efficient cpSRP-dependent LHCP targeting to the thylakoid membrane depends on complex formation between cpSRP43 and cpSRP54 *in vivo*. Since our data suggest that free cpSRP54 does not support LHCP insertion, it will be interesting to analyse the transport of LHCP proteins in the cpSRP43-knockout mutant *chaos*, which shows reduced but still considerable amounts of LHCPs (Amin et al., 1999; Klimyuk et al., 1999).

Our findings substantiate the hypothesis that the evolution of the C-terminal tail region was likely triggered to facilitate efficient posttranslational sorting of the LHCP proteins in land plants (Dünschede et al., 2015). Consistent with this, our data demonstrate that the plant-specific C-terminal region of cpSRP54 is dispensible for the cotranslational transport of the photosynthetic reaction center subunits, while the cpSRP54 G and M domain, which are conserved in all SRP systems, are important for cotranslational sorting. Binding of cpSRP54 to ribosomes was shown to depend on two binding motifs in the M domain and the C-terminal tail region (Hristou et al., 2019). Accordingly, we did not observe binding of the cpSRP54 variant lacking these regions to ribosomes, while the variant without the C-terminal tail showed still some binding (Supplementary Figure S2). Interestingly, the M domain of the chloroplast SRP system plays also an important role in the docking of the ribosome nascent chain complex to the thylakoid membrane as it greatly accelerates the binding to its receptor, cpFtsY, a function that is mediated by the SRP RNA in the bacterial system (Jaru-Ampornpan et al., 2009). In analogy to the bacterial SRP system, it can be speculated that the M domain is additionally able to bind directly to the nascent chain, while it emerges out of the ribosomal peptide tunnel exit site. A direct contact between cpSRP54 and the nascent chain of D1 was already demonstrated, but molecular details of this interaction are currently unclear (Nilsson et al., 1999; Nilsson and van Wijk, 2002).

The complex formation of the SRP GTPases cpSRP54/cpFtsY is GTP-dependent as binding of the nucleotide leads to conformational changes of the proteins enabling dimerization and therefore reciprocal GTPase activation (Jaru-Ampornpan et al., 2007; Nguyen et al., 2011; Wild et al., 2016). While *in vitro* experiments demonstrated that LHCP insertion into thylakoid membranes is indeed GTP-dependent (Hoffman and Franklin, 1994; Yuan et al., 2002), the importance of GTP hydrolysis for cotranslational targeting in chloroplasts remained unknown. Here, we observed that a point mutation within the highly conserved GI motif of cpSRP54 at amino acid position 185 dramatically affects the functionality of cpSRP54 in post– and cotranslational sorting of LHCPs and plastid-encoded photosynthetic reaction center proteins, respectively, *in vivo*. The GI motif, also known as Walker-A motif or P-loop, is part of the active center for GTP binding/hydrolysis and interacts with the β-phosphate of GTP by forming an anion hole (Shan, 2016; Wild et al., 2016). Therefore, it seems reasonable that mutations within the GI motif can affect the nucleotide affinity and GTP hydrolysis rate as we observed for the cpSRP54(Q185R) variant.

Our investigation of thylakoid membrane complexes revealed that an intermediate assembly state of PSI (PSI*) is slightly upregulated in *ffc* and greatly accumulates in cpSRP54ΔC/*ffc*. This complex is composed of a subset of PSI core proteins and lacks the LHCI antenna proteins. Therefore, it resembles previously described PSI assembly intermediates in *N. tabacum*, *Chlamydomonas reinhardtii* and a recently identified rice mutant lacking cpSRP54 (Ozawa et al., 2010; Wittenberg et al., 2017; Gao et al., 2022). The specific accumulation of this assembly complex in the *ffc* mutant expressing cpSRP54 lacking its C-terminal region can be explained by our findings that this cpSRP54 variant still supports the cotranslational biogenesis of the PSI core proteins PsaA and PsaB, while the posttranslational transport of the LHCI antenna proteins is impaired. Currently it is known that PSI assembly in land plants begins with the formation of the PsaA/PsaB heterodimer and continues with the integration of a specific subset of peripheral subunits (PsaC-E, PsaH, PsaI and PsaL) to form PSI*, a stable PSI core assembly intermediate. Subsequently, the addition of PsaF is required to trigger full assembly of PSI by the integration of five additional subunits, including PsaK, and the LHCI proteins (Schöttler et al., 2011; Wittenberg et al., 2017; Zhang et al., 2024). Notably, we observed that the reduced assembly of LHCI to PSI leads to a reduced amount of PsaK in PSI. In this respect, it is important to note that PsaK was shown to integrate into the thylakoid membrane independently of cpSRP54 (Mant et al., 2001). Furthermore, the cpSRP54ΔC/*ffc* plant shows a specific down-regulation of PsaK in PSI but no changes in the total amount of PsaK demonstrating that PsaK insertion itself is not impaired in cpSRP54ΔC/*ffc*. Therefore, our data indicate that the integration of PsaK in PSI needs the assembly of LHCI and presents one of the last steps in PSI maturation in land plants.

## Supplementary data

The following supplementary data are available at JXB online.

### Supplementary methods

Sucrose density gradient centrifugation. Genomic DNA extraction and genotyping PCR. Size exclusion chromatography.

**Table S1:** Primer used for generation of transformation constructs, site-directed mutagenesis, and genotyping PCRs.

**Table S2:** Isothermal titration calorimetry (ITC) parameters – Thermodynamic values determined from the ITC experiments.

**Figure S1:** Identification of *A. thaliana ffc*-complementation lines.

**Figure S2:** Sucrose density gradient analysis of stromal extracts from *A. thaliana* wild type (WT), *ffc* and *ffc*-complementation lines.

**Figure S3:** Two-dimensional BN-PAGE/SDS-PAGE and immunoblot analysis of thylakoid membrane multiprotein complexes in *A. thaliana ffc*.

**Figure S4:** Analysis of the migration behaviour of recombinant cpSRP54 and the point mutation variant cpSRP54(Q185R).

## Abbreviations

(cpSRP): chloroplast signal recognition particle
(LHCPs): light-harvesting chlorophyll a/b-binding proteins
(PSI): photosystem I
(PSII): photosystem II
(LHCI): light-harvesting antenna complexes I
(LHCII): light-harvesting antenna complexes II

## Acknowledgements

We like to acknowledge Silke Funke for excellent technical assistance. We thank Annika Ehlert, Julia Bollmann and Laura Czech for help with the experiments. This work was supported by the Deutsche Forschungsgemeinschaft (SCHU 1163/ 6-2 to D.S.). G.B. acknowledges support by the ERC AdV grant KIWIsone (Grant agreement ID: 101019765).

## Author contribution

A.B., J.O., B.D., G.B. and D.S. conceived the idea and designed the research. A.B. and J.O. implemented the *in vivo* experiments and executed the biochemical and statistical analyses. V.Z. and P.B. performed the *in vitro* ITC and GTPase activity measurements. A.B. and D.S. wrote the article.

